# Stimulus-related modulation in the 1/f spectral slope suggests an impaired inhibition of irrelevant information in people with multiple sclerosis

**DOI:** 10.1101/2023.12.28.573572

**Authors:** Fahimeh Akbarian, Chiara Rossi, Lars Costers, Marie B D’hooghe, Miguel D’haeseleer, Guy Nagels, Jeroen Van Schependom

## Abstract

**Background:** Multiple sclerosis (MS) is an inflammatory and neurodegenerative disease characterized by neuronal and synaptic loss, resulting in an imbalance of excitatory and inhibitory synaptic transmission. MS leads to cognitive impairment such as reduced information processing speed and impaired working memory (WM). Recent studies have suggested that the 1/f slope of EEG/MEG power spectra can be associated with the excitation/inhibition (E/I) balance. A normal E/I balance is crucial for normal information processing and working memory.

**Methods:** We analyzed magnetoencephalographic (MEG) recordings of 38 healthy control subjects and 79 people with multiple sclerosis (pwMS) while performing an n-back working task. We computed and compared the steepness of the 1/f spectral slope through the FOOOF algorithm in the time windows [-1 0] and [0 1] s peristimulus time for both target and distractor stimulus for each brain parcel and for different working memory loads (0-back, 1-back, 2-back).

**Results:** The spectral slope was significantly steeper after the stimulus onset and was correlated with reaction time. We also observed a steeper 1/f slope after distractor stimuli in healthy subjects compared to pwMS. Finally, we observed significant correlations between the 1/f spectral slope modulation and visuospatial working memory functioning in both healthy subjects and pwMS.

**Conclusion:** Our findings are consistent with an increased inhibition following stimulus onset. In pwMS, this increase is reduced, suggesting dysfunctional inhibition of irrelevant information. Finally, this impaired modulation is significantly associated with a pencil-paper test of visuospatial working memory.

**Highlights:**

- The flatter 1/f slope after distractor stimuli in people with multiple sclerosis (pwMS) compared to healthy subjects suggests a less pronounced inhibition of irrelevant information in pwMS.
- The significantly flatter 1/f slope was observed in the left inferior dorsal prefrontal cortex of pwMS in both 1-back and 2-back conditions. This particular brain parcel is known for its key role in motor planning, and the maintenance of sustained attention and working memory and executive functions.
- A steeper 1/f slope after target and distractor stimuli suggests an increase in inhibition following stimulus onset in both healthy controls (HCs) and people with MS (pwMS).
- The 1/f slope modulation correlates with visuospatial working memory performance.

## 1. Introduction

Multiple sclerosis (MS), an inflammatory and neurodegenerative disease characterized by inhibitory and excitatory synaptic loss (Di Filippo et al., 2018; Huiskamp et al., 2022; Mandolesi et al., 2015), frequently imposes a significant cognitive burden on people with MS (pwMS). Cognitive domains such as information processing speed (DeLuca et al., 2004; Van Schependom et al., 2015) and working memory (Chiaravalloti and DeLuca, 2008; D’Esposito et al., 1996) are significantly affected.

Working memory (WM) is the capacity to actively and temporarily retain and manipulate a limited amount of information for a brief period of time (Baddeley, 2012). Working memory is crucial for a variety of cognitive processes, including learning, problem solving, decision making, and comprehension (Baddeley, 2003). It has been computationally demonstrated that the balance between excitation and inhibition (E/I) is essential for efficient information processing and working memory maintenance (Lim and Goldman, 2013; Vogels and Abbott, 2009). In addition, the suppression of irrelevant information besides the selection and storage of relevant information seems to play an important role in working memory processes (Getzmann et al., 2018). Evidence suggests that better inhibition of distractors is associated with improved retrieval of task-relevant information (Bonnefond and Jensen, 2012; Griffin and Nobre, 2003; Matsukura et al., 2007; Schneider et al., 2017).

The use of magneto/electroencephalography (M/EEG) in the frequency domain has been extensively employed in working memory studies to investigate oscillatory modulations. Previous research has e.g., shown alpha suppression during the encoding and delay period of visual stimuli (Gevins, 1997; Sauseng et al., 2009). Nevertheless, it has also been shown that the non-oscillatory component of neuronal activity contains functionally relevant information. The non-oscillatory or 1/f-like shape of the power spectrum of neuronal activity refers to a power decrease as frequency increases. The slope, representing the steepness of this 1/f power-law component (denoted as "*x*" in 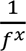), has been associated with the excitation/inhibition ratio (Gao et al., 2017), where a steeper slope indicates a higher level of inhibition (Akbarian et al., 2023; Chini et al., 2022; Gao et al., 2017).

A recent study (Virtue-Griffiths et al., 2022) demonstrated that the task-related 1/f slope modulation was independent of alpha suppression. This modulation was region-specific, with a steeper slope observed in the fronto-central area and a flatter slope in the lateral parieto-occipital region. Moreover, Gyurkovics et al (Gyurkovics et al., 2022) were the first to reveal an immediate and possibly transient increase in 1/f slope following an oddball auditory stimulus. This finding suggests a global increase in inhibitory activity and implies a momentary interruption of ongoing processing to facilitate further stimulus-specific processing.

In the current study, we use a visual-verbal n-back task to examine the modulation in the 1/f slope. We hypothesize that the 1/f slope will be steeper post- vs. pre-stimulus during the n-back task, similar to what has been observed in recent studies (Gyurkovics et al., 2022; Virtue-Griffiths et al., 2022). We also hypothesize a correlation between flatter 1/f slope and longer reaction times, as shown by a recent study (Kałamała et al., 2023) where individuals with flatter 1/f slope have a higher inverse efficiency score (i.e., adjusted reaction times). In addition, we hypothesize that an impaired excitation/inhibition balance – as suggested in pwMS (Huiskamp et al., 2022) – will lead to reduced inhibition during distractor trials. Finally, we hypothesize that the level of 1/f modulation throughout a cognitive task is associated with offline cognitive performance.

## 2. Methods

### 2.1. Participants

MEG data were acquired in 117 subjects during the n-back task: 38 healthy subjects and 79 people with multiple sclerosis (pwMS). Since in our recent study (Akbarian et al., 2023) we observed that benzodiazepines affect the 1/f spectral slope, we divided the group of pwMS into two separate groups based on their treatment: 19 pwMS treated with benzodiazepines (pwMS(BZDp)) and 60 pwMS not treated with benzodiazepines (pwMS(BZDn)). All pwMS were recruited at the National MS Center Melsbroek and were required to meet the diagnostic criteria for multiple sclerosis according to McDonald’s revised criteria (Polman et al., 2011) while having an Expanded Disability Status Scale (EDSS) score (Kurtzke, 1983) of less than or equal to 6. Exclusion criteria included experiencing a relapse or receiving corticosteroid treatment within the preceding six weeks, as well as having a pacemaker, dental wires, major psychiatric disorders, or epilepsy. In Table 1, we present a detailed description of people with MS and healthy controls.

**Table 1.** Description of subjects. We report the mean values and standard deviations for different clinical parameters. For EDSS the median and interquartile range (IQR) are shown. The comparisons were performed using permutation testing with N = 5000 for all parameters except gender for which a chi-squared test was used. HC: healthy control, pwMS(BZDp) and pwMS(BZDn): pwMS with and without benzodiazepines.

|  | HC | pwMS sub-groups |  | Group differences (p-values) |  |  |
| --- | --- | --- | --- | --- | --- | --- |
|  | HC | pwMS(BZDp) | pwMS(BZDn) | HC vs pwMS(BZDn) | HC vs pwMS(BZDp) | pwMS(BZDn) vs pwMS(BZDp) |
| *N* | 38 | 19 | 60 | - | - | - |
| *Gender (M/F)* | 15/23 | 0/19 | 22/38 | 0.94 | 0.0040 | 0.0048 |
| *Age (yrs)* | 47±11 | 47±7 | 48±10 | 0.89 | 0.99 | 0.91 |
| *Education (yrs)* | 15.2±2 | 13±2 | 14±2 | 0.16 | 0.001 | 0.08 |
| *Disease duration(yrs)* | - | 13±6 | 15±8 | - | - | 0.34 |
| *EDSS median [IQR]* | - | 3.5 [3-4.25] | 2.5 [2-3.5] | - | - | 0.051 |
| *Cognitive assessment* |  |  |  |  |  |  |
| *SDMT* | 53 ±9 | 47±7 | 49±12 | 0.07 | 0.01 | 0.49 |
| *VGLT* | 66 ±7 | 62±10 | 63±10 | 0.29 | 0.12 | 0.56 |
| *COWAT* | 17±4 | 17±4 | 15±5 | 0.045 | 0.65 | 0.33 |
| *BVMT-R* | 28±5 | 24 ±6 | 26±6 | 0.054 | 0.03 | 0.57 |

### 2.2. MEG data acquisition

The MEG acquisition for the first 30 pwMS and 14 healthy subjects was done using an Elekta Neuromag Vectorview scanner (Elekta Oy, Helsinki, Finland) and the rest of the subjects were scanned using an upgraded scanner, the Elekta Neuromag Triux scanner (MEGIN, Croton Healthcare, Helsinki, Finland) due to a system upgrade at the CUB Hôpital Erasme (Brussels, Belgium). Both MEG scanners used a sensor layout as 102 triple sensors, each consisting of one magnetometer and two orthogonal planar gradiometers and were placed in a lightweight magnetically shielded room (MaxshieldTM, Elekta Oy, Helsinki, Finland). Prior to MEG acquisition, four head position indicator coils were attached to the left and right mastoid and forehead of the subject to track the head position during the data acquisition. The location of these coils and at least 400 head-surface points (on the nose, face, and scalp) with respect to anatomical fiducials (nasion, left and right preauricular) were determined with an electromagnetic tracker (Fastrak, Polhemus, Colchester, Vermont, USA). MEG data were captured using a sampling frequency of 1,000 Hz and a band-pass filter ranging from 0.1 Hz to 330 Hz. Participants were asked to maintain maximum stillness throughout the data acquisition process. They were seated upright, with their heads positioned towards the back of the MEG helmet. An electrooculogram (EOG) and electrocardiogram (ECG) were simultaneously recorded with MEG signal acquisition to be used in offline artefact rejection.

### 2.3. N-back task

During the MEG acquisition, all participants were asked to perform an n-back task (Costers et al., 2020) with three conditions or levels of WM load (0, 1 and 2-back). The examiner instructed participants to press a button with their right hand when the letter displayed on the screen was the letter X (0-back condition), the same letter as the one before (1-back condition), or the same letter as two letters before (2-back condition). Figure 1 demonstrates the n-back paradigm. The 6 x 6.5 cm letter stimuli were projected onto a screen 72 cm from the front of the MEG helmet. Each stimulus was displayed on the screen for 1 second, with an inter-trial interval of 2.8 seconds. The instructions for the specific conditions were presented at the start of every block for 15 seconds. Using a photodiode, the onset of the visual stimuli was measured. The time between the photodiode detecting stimulus onset and the subject pressing the button was then calculated as the reaction time.

**Figure 1.**
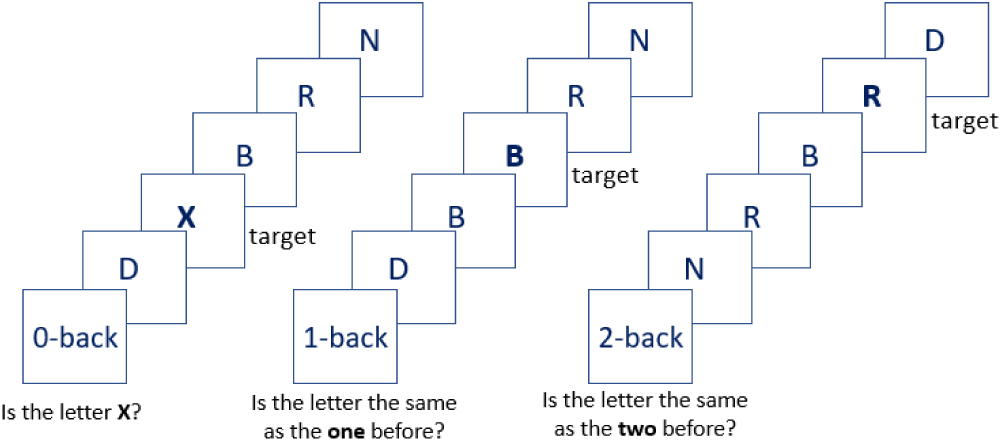
Illustration of the visual-verbal n-back paradigm. The stimulus duration was 1 second with intervals of 2.8 seconds (Costers et al., 2020).

The task was divided into four blocks per level, with a total of 12 blocks presented in a random sequence. Each block contained 20 stimuli, for a total of 240 stimuli and 25, 23, and 28 target trials per condition, respectively. Importantly, we also included non-target (distractor) trials in our analysis, and we excluded trials in which an incorrect button press was recorded.

### 2.4. MEG processing and parcellation

First, we used the temporal extension of the signal space separation algorithm (MaxfilterTM, Elekta Oy, Helsinki, Finland, version 2.2 with default parameters) to remove the external interferences and correct head movements (Taulu et al., 2005). Then the preprocessing steps were done using Oxford’s Software Library (OSL) pipeline built upon FSL, SPM12 (Welcome Trust Centre for Neuroimaging, University College London) and Fieldtrip (Oostenveld et al., 2011). We first applied an FIR antialiasing lowpass filter, then we downsampled the data to 250 Hz. Using OSL’s RHINO (https://github.com/OHBA-analysis) algorithm, data were automatically coregistered with the subject’s T1 image: the head shape points were coregistered to the scalp extracted using FSL’s BETSURF and FLIRT (Jenikson, Mark, 2005; Smith, 2002) and then transformed into a common MNI152-space (Mazziotta et al., 1995). Data were filtered between 0.1 and 70 Hz and then a 5^th^ order Butterworth filter (between 49.5 and 50.5 Hz) was applied to remove the power line noise.

A semi-automated independent component analysis (ICA) algorithm was employed to visually identify and remove ocular and cardiac artefacts based on the correlation of the components’ time series with EOG and ECG signals respectively. Next, we used a linearly constrained minimum variance (LCMV) beamformer to project the MEG data to a source space (Oostenveld et al., 2011; Quinn et al., 2018; Woolrich et al., 2011). The source-reconstructed data were then parceled using a parcellation atlas including 42 parcels as published before (Vidaurre et al., 2018). The first principal component (PC) of the different time series within each parcel was used as the time series for that parcel. As previously discussed, the parcellation atlas covered the entire cortex and did not include subcortical areas (Van Schependom et al., 2019; Vidaurre et al., 2018).

### 2.5. Power spectral analysis and estimation of the aperiodic components

For each trial, we analyzed a time window spanning one second both before and after the onset of stimuli. Within this timeframe, we computed the power spectral density (PSD) using the default Welch function in SciPy with a window length of 250 samples (one second) at each brain parcel for every subject. Then for each time window before and after stimuli, we averaged the power spectra over trials within each subject. Further, we applied the "fitting oscillations and one over f" (FOOOF) algorithm, as introduced by (Donoghue et al., 2020), to estimate the 1/f exponent for the resulting power spectra. We defined the fitting frequency range between 3 Hz to 45 Hz. We set the lower border of the fitting frequency range to 3 Hz to avoid the aperiodic differences driven by delta oscillations and the upper border to 45 Hz to avoid the notch-filtered peak around 50 Hz due to power line noise. Since the power spectrum followed an almost linear line in log-log space within the selected frequency range, we used the “fixed” mode for parameterization.

The 1/f slope refers to the steepness of the 1/f aperiodic component (exponent "*x*" in 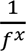). We used the “1/f slope change” term when comparing the pre- versus post-stimulus 1/f slope at a group level and we defined the “1/f slope modulation” term as the post-stimulus slope minus the pre-stimulus slope within each subject similar to (Gyurkovics et al., 2022).

### 2.6. Periodic alpha power estimation

To estimate the periodic alpha power, we first separated the periodic component by correcting the power spectrum density for the aperiodic component fitted. After subtraction of the 1/f component, the periodic alpha power was defined as the average alpha power in the 8 Hz to12 Hz band.

### 2.7. Neuropsychological assessment

Neuropsychological tests were performed on the same day as the MEG recording for all subjects. The neuropsychological tests included the Symbol Digit Modalities Test (SDMT, (A Smith, 1968)) to capture information processing speed, the Dutch version of the California Verbal Learning Test (CVLT-II, (Strober et al., 2009)), Dutch version: VGLT to assess verbal memory, the Controlled Oral Word Association Test (COWAT, (Ruff et al., 1996)) to assess verbal fluency and the Brief Visuospatial Memory Test (Revised; BVMT-R, (Benedict et al., 1996)) to assess spatial memory. Fatigue is assessed by the Fatigue Scale for Motor and Cognitive Function (FSMC, (Penner et al., 2009)) and depression by Beck’s Depression Inventory (BDI, (Beck, 1961)).

### 2.8. Statistics

All statistical analyses used non-parametric measures, which do not rely on assumptions about the distribution of the data. The Wilcoxon signed-rank test was performed for the paired-wise comparisons. For the non-paired comparisons, we used either the Wilcoxon rank-sum test or the Mann-Whitney U test depending on the tie presence (i.e., two or more differences have the same absolute value) between two samples. When calculating tests for multiple cortical parcels, the Benjamini-Yekutieli false discovery rate (FDR, (Benjamini and Hochberg, 1995)) correction for multiple comparisons was applied to the p-values. We used the “cocor” package (Diedenhofen and Musch, 2015) to compare the dependent and independent correlations. For all p-values, the cutoff of 0.05 was used.

### 2.9. Ethics

The research was approved by the University Hospital Brussels’s local ethics committees of the University Hospital Brussels (Commissie Medische Ethiek UZ Brussel, B.U.N. 143201423263, 2015/11) and the National MS Center Melsbroek (2015-02-12). All participants also provided written informed consent.

**Figure 2.**
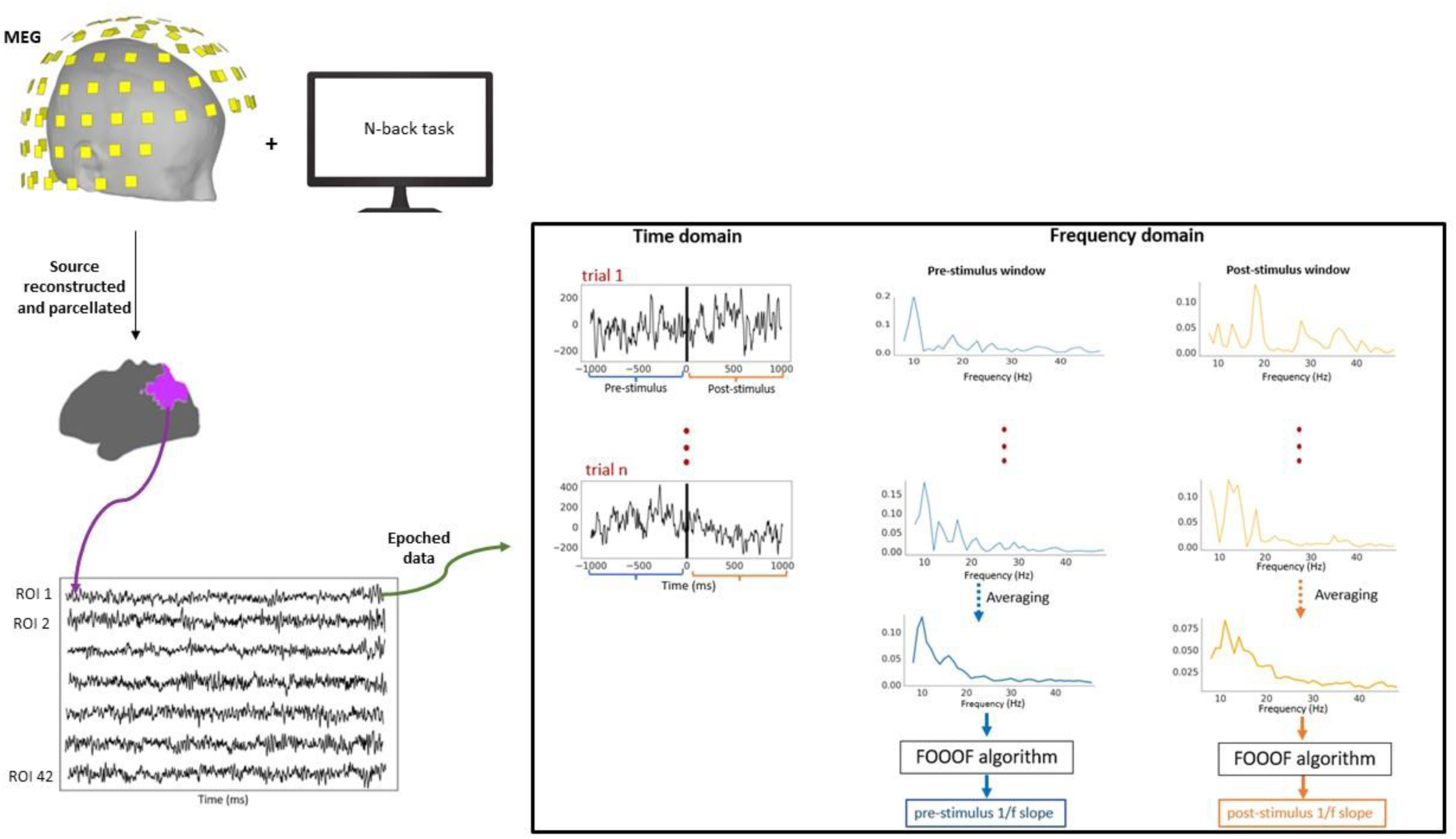
An illustration of the data analysis for a single subject, in a single condition, for a single trial and at a single brain parcel. We determined the time window of one second before and after the onset of stimuli. Within this timeframe, we computed the PSD for each trial at each brain parcel for every subject. Then, for each time window before and after stimuli, we averaged the power spectra over trials within each brain parcel for each subject. Further, we applied the "fitting oscillations and one over f" (FOOOF) algorithm (Donoghue et al., 2020) to estimate the 1/f exponent for the resulting power spectra.

## 3. Results

### 3.1. Behavioral data

Figure 3 shows an increasing median reaction time with increasing WM load (0, 1, 2-back conditions) within each group of subjects. The median RT during the 0-back is significantly faster than the 2-back (p<0.0001) and the same holds for the 1-back as compared to the 2-back (p<0.007). These results are in line with WM literature in which the direct positive association between reaction times and WM load is frequently reported. Further, we compared the reaction time (RT) and accuracy between healthy subjects, pwMS(BZDn) and pwMS(BZDp). We observed a significantly longer reaction time in pwMS(BZDn) and pwMS(BZDp) compared to the HC group. We also observed a significant difference in response accuracy between healthy subjects and pwMS(BZDp) (p < 0.0001) in the 1-back condition and between healthy subjects and pwMS(BZDn) (p = 0.043) in the 2-back condition, with better performance in HC subjects (Supplementary Figure S1).

**Figure 3.**
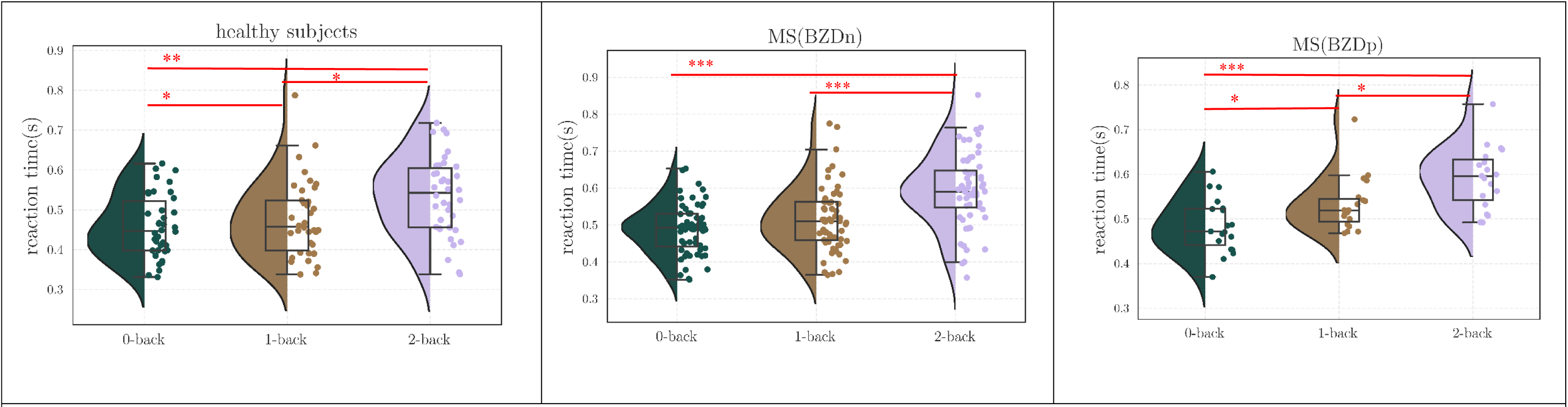
The distribution of reaction times for three groups (HCs, pwMS(BZDn), pwMD(BZDp)) and three conditions (0-back, 1-back, 2-back). The Wilcoxon signed-rank test was used to compare the reaction times between different conditions within each group of study. *:0.001<p<0.05, **: 0.0001<p<0.001, ***: 0.00001<p<0.0001.

### 3.2. Stimulus-related changes in the 1/f slope

We compared the 1/f slope pre vs post stimulus. This analysis was performed separately for target and distractor stimuli.

#### 3.2.1. Steeper 1/f slope in post-stimulus target trials

In the case of target stimuli, we observed a statistically significant increase in the 1/f slope following stimulus onset across all three conditions (0-back, 1-back, and 2-back) for both healthy subjects and pwMS(BZDn). However, this increase in the 1/f slope was not statistically significant for pwMS(BZDp), neither when considering the entire brain nor at parcel level.

Figure 4A shows the averaged 1/f slope pre- and post-stimulus in the 0-back condition. Figure 4B presents the same analysis but on a parcel level. While the results for the 0-back condition of pwMS(BZDn) are displayed here, the results for the other cohorts are very similar (see Supplementary Figure S2).

**Figure 4.**
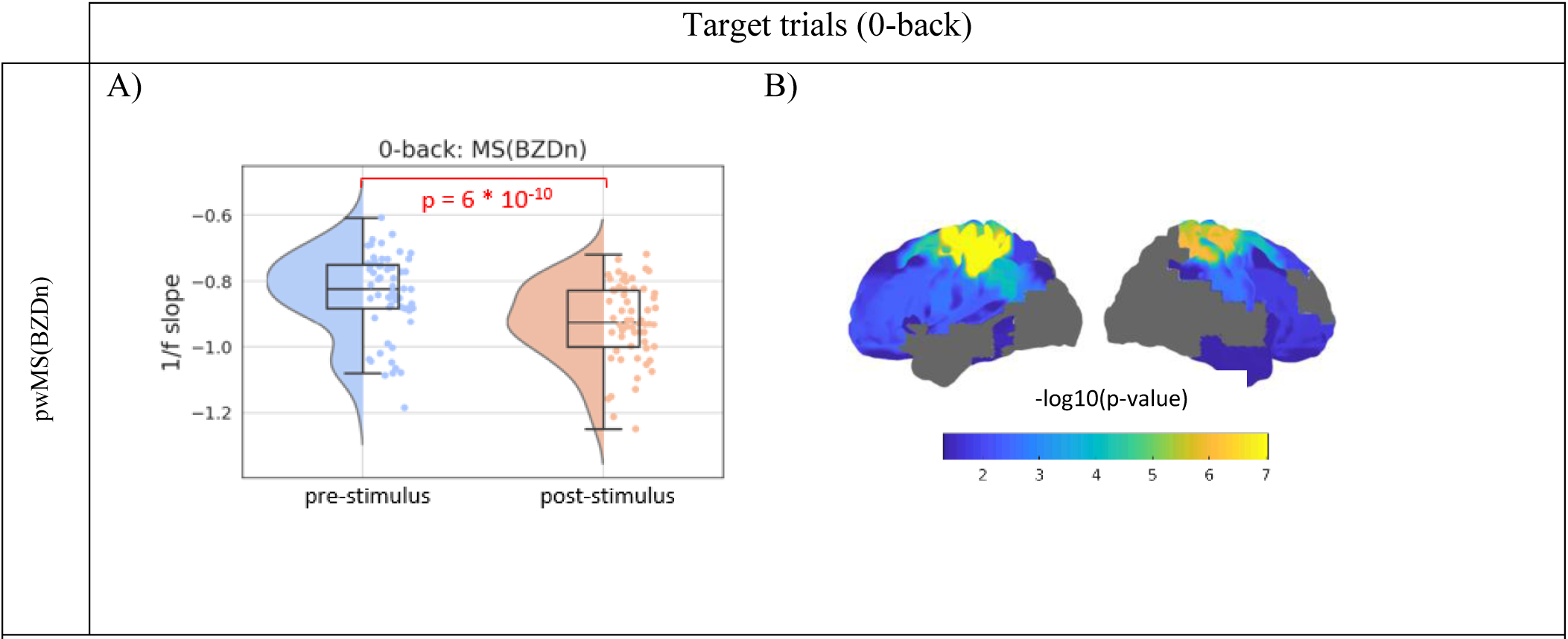
The pre- vs post-stimuli 1/f slope for pwMS(BZDn) group in target trials and 0-back condition, for 1/f slope **A)** averaged over the whole brain and **B)** in parcel level. Note that the -log10(p-value)>1.3 indicated the significant values and that p-values have been corrected for multiple comparisons across the 42 parcels. Wilcoxon signed-rank test was used to compare the pre- and post-stimuli 1/f slope.

#### 3.2.2. Steeper 1/f slope in post-stimulus distractor trials

Similar to the analysis of the target stimuli, we observe a significantly steeper slope after the distractor stimuli, suggesting that the observed alteration in the 1/f slope cannot be solely attributed to the act of button pressing. Figure 5 provides a visual representation of the averaged 1/f slope changes over the entire brain before and after the onset of distractor stimuli along with the cortical distribution of regions displaying statistically significant variations in 1/f slope between the pre- and post-stimulus onset. The results of all three groups of subjects for the three 0-back, 1-back and 2-back conditions are shown. The reported p-values have been corrected for multiple comparisons. The most significant results are seen in the pwMS(BZDn) group. However, it should be noted that this is also the largest group, and the effect sizes are similar across all comparisons.

**Figure 5.**
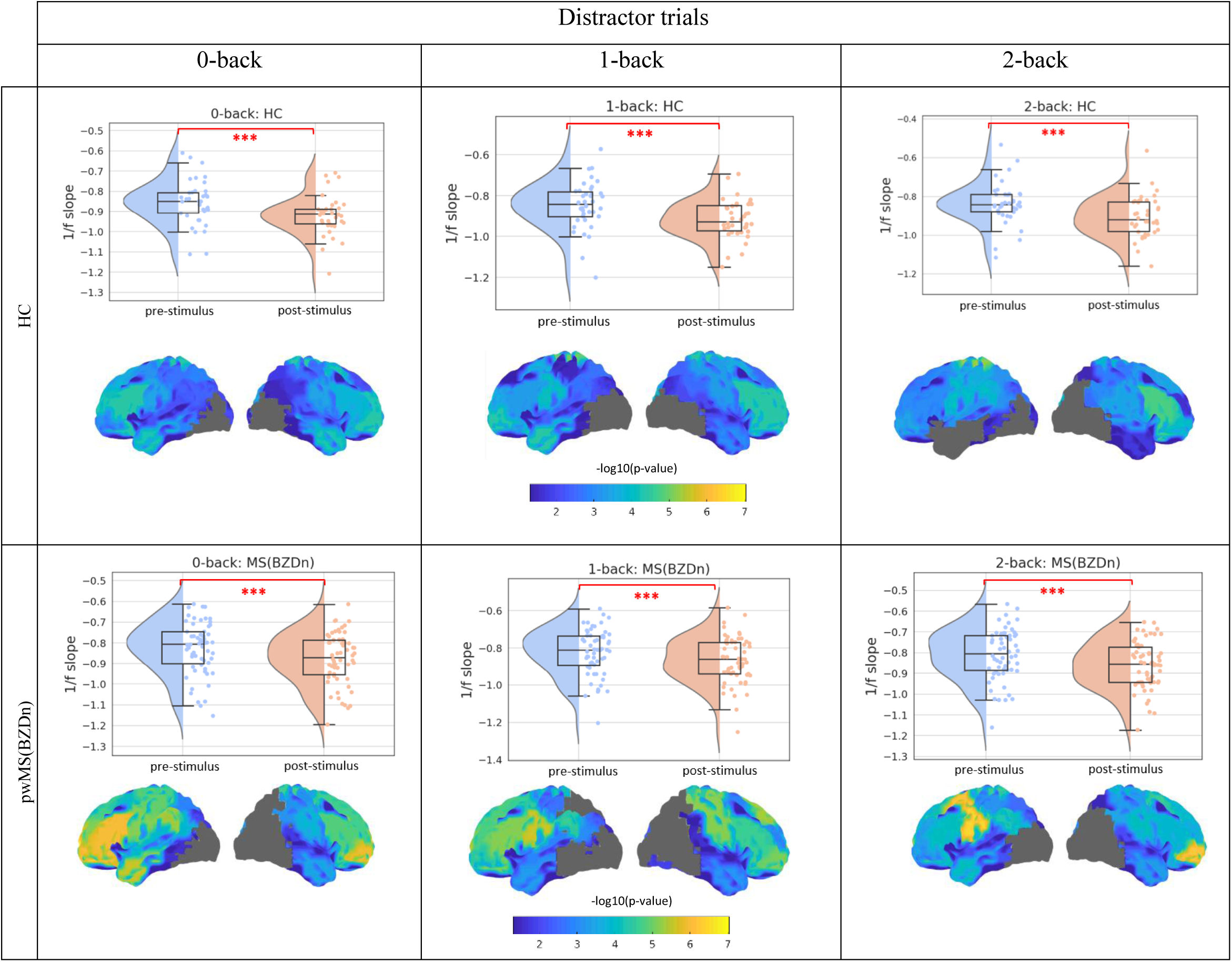
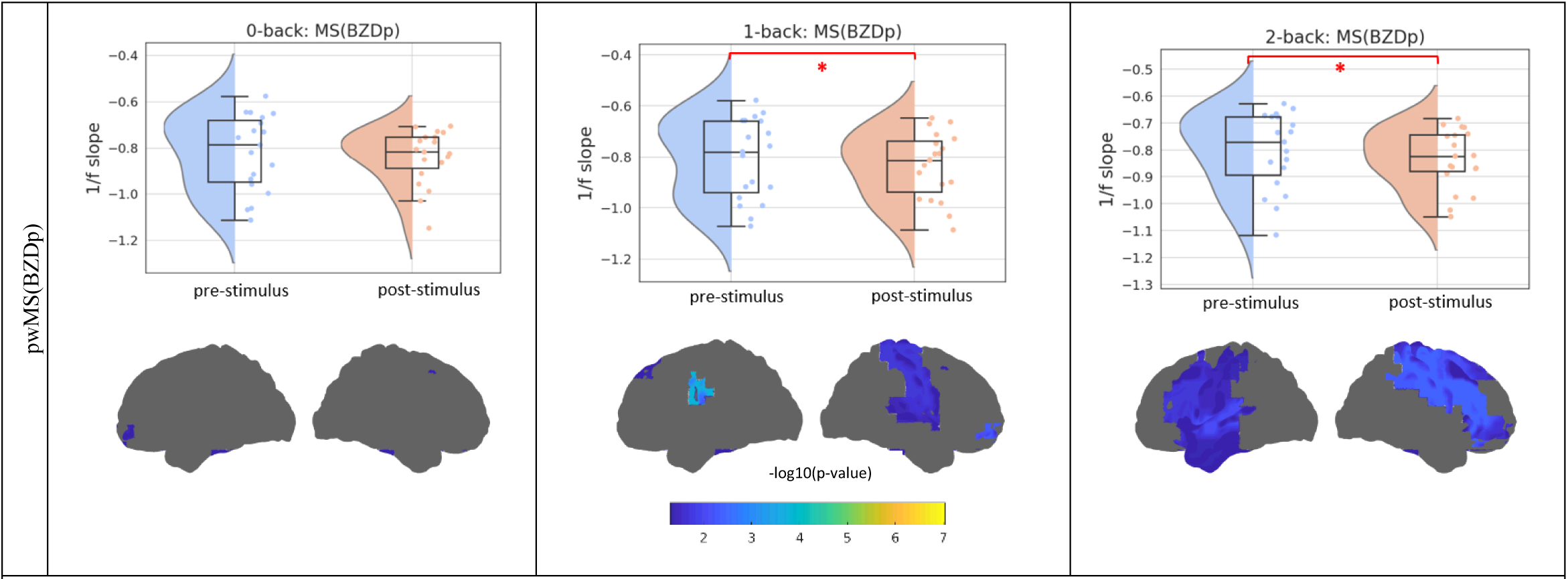
The pre- vs post-stimuli 1/f slope in distractor trials and three conditions (0-back, 1-back and 2-back), for both 1/f slope averaged over the whole brain and in parcel level. Note that the -log10(p-value)>1.3 indicated the significant values and that p-values have been corrected for multiple comparisons across the 42 parcels. The Wilcoxon signed-rank test was used to compare the pre- and post-stimuli 1/f slope. First row: HC subjects, second row: pwMS(BZDn), third row: pwMS(BZDp). *:0.001<p<0.05, **: 0.0001<p<0.001, ***: 0.00001<p<0.0001.

### 3.3. Comparing the 1/f slope between HC and pwMS

We compared the 1/f slope in target and distractor stimuli between healthy subjects and pwMS(BZDn) and pwMS(BZDp). After distractor stimuli, healthy subjects demonstrated a significantly steeper slope compared to the pwMS(BZDn) and pwMS(BZDp) in all three conditions. Importantly, the 1/f slope before and after target stimuli and before distractor stimuli was not significantly different between HC and pwMS(BZDn). As it is shown in Figure 6, there was no significant difference in 1/f slope between pwMS(BZDn) and pwMS(BZDp) after distractor trials.

**Figure 6.**
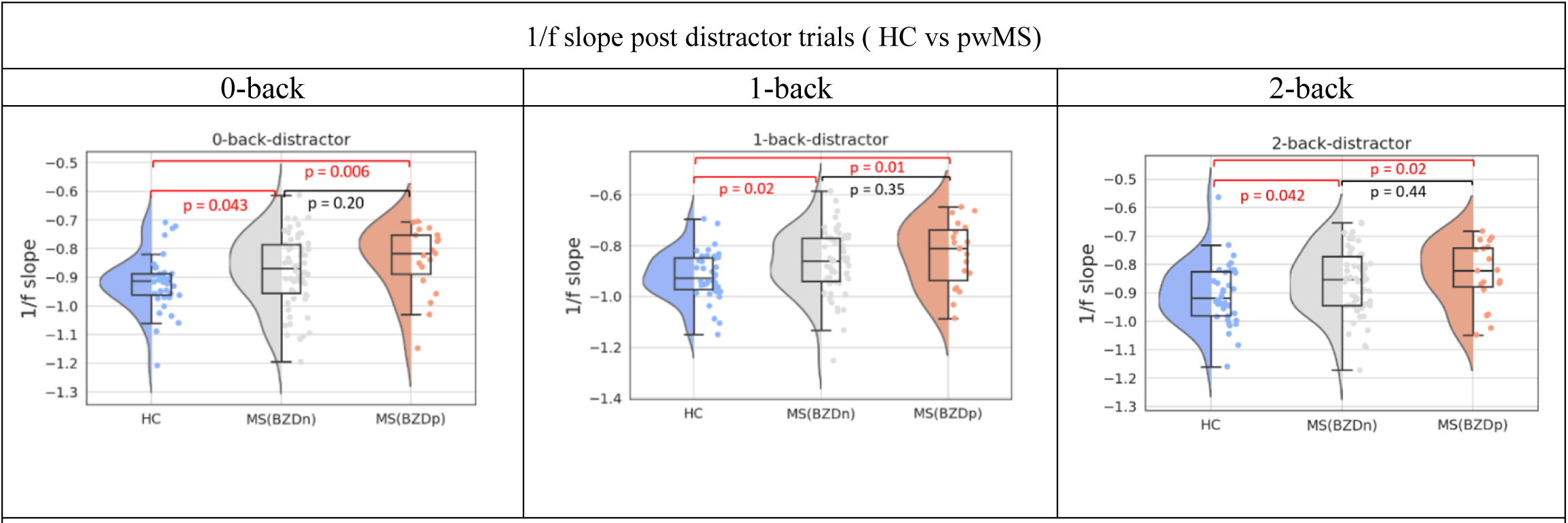
The distribution of post-distractor stimuli 1/f slope -averaged over the whole brain- for the HC and pwMS groups in all three conditions (0-back, 1-back and 2-back). The Wilcoxon rank-sum test was used.

### 3.4. Correlation between 1/f slope pre- and post-stimulus and reaction time

Further, we calculated the correlation between the subjects’ mean reaction time and their 1/f slope both before and after the onset of target and distractor stimuli across all subjects. This analysis was performed across all three cognitive load conditions: 0-back, 1-back, and 2-back, see Figure 7 for target trials and supplementary Figure S5 for distractor trials. As expected, our results revealed that a flatter slope (less inhibition) is linked to a longer reaction time. Notably, this correlation was more pronounced in the 1-back and 2-back conditions compared to the 0-back condition indicating a stronger relationship in the context of a higher cognitive load.

**Figure 7.**
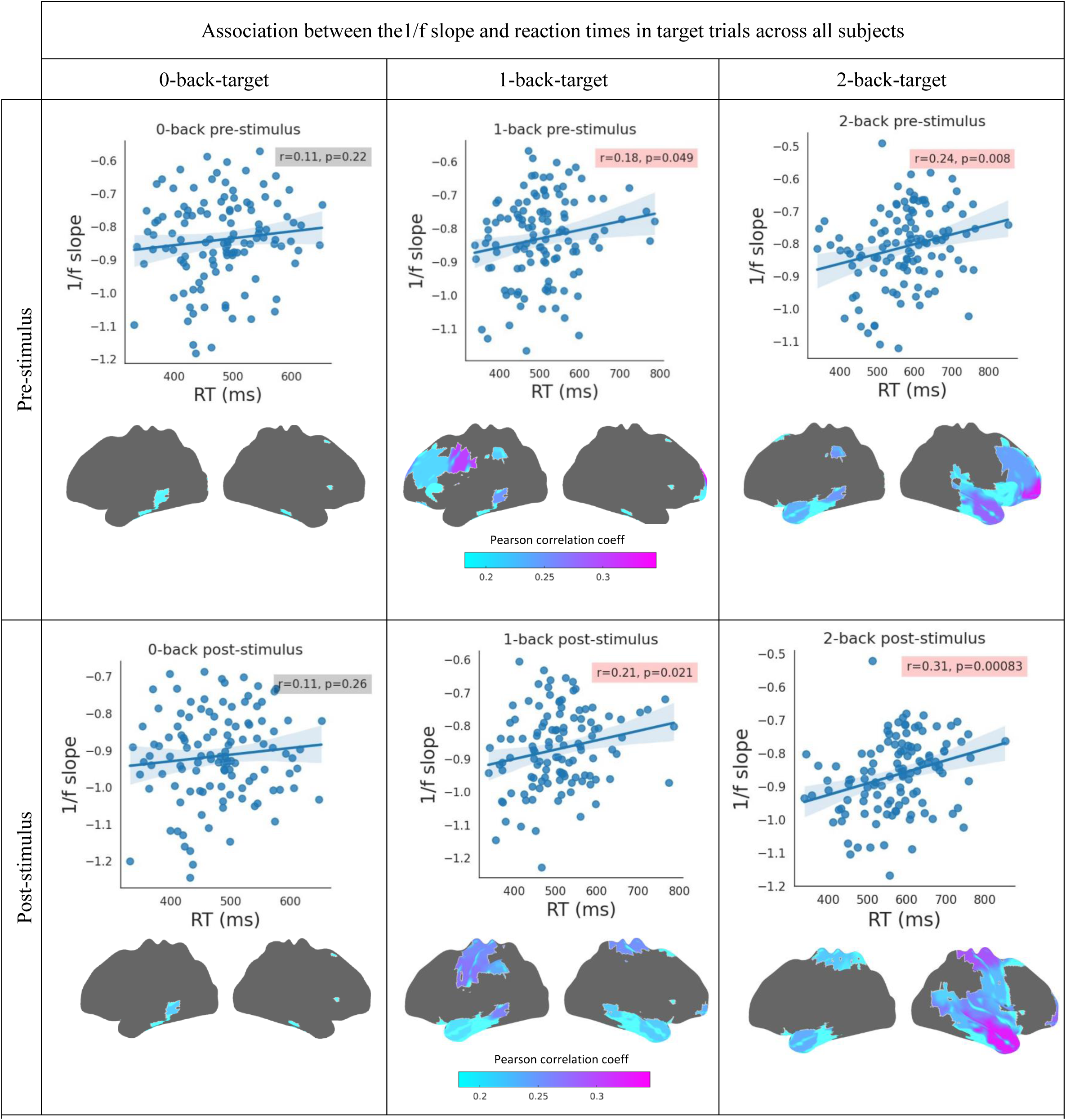
Correlation analysis between reaction time and 1/f slope of the time window one second pre- and one second post-target stimulus onset both for averaged over the whole brain and in parcel level over all subjects. In each grid, the scatter plot shows the correlation for the averaged 1/f slope over the whole brain and reaction times and the cortical map shows the parcels with significant correlation for the parcel level analysis. We included all subjects to increase the statistical power.

### 3.5. The relationship between the 1/f slope modulation and cognitive performance

To investigate the potential association between the 1/f slope modulation and cognitive performance, we conducted a Pearson correlation analysis between the 1/f slope modulation induced by stimuli and the offline cognitive performance measured through standardized neuropsychological tests. Our analysis focused exclusively on healthy subjects and pwMS who did not receive benzodiazepines, as the cohort of pwMS who had received benzodiazepines (pwMS(BZDp)) did not exhibit significant differences in task-induced 1/f slope modulation compared with the other groups.

Our results revealed a significant positive correlation between the 1/f slope modulation and BVMT scores as a measure of visuospatial working memory performance among healthy subjects. Healthy subjects with better visuospatial working memory performance tend to have a smaller increase in inhibition post-stimulus. Conversely, a significant negative correlation was observed between these variables among pwMS who had not received benzodiazepines (pwMS(BZDn)), see Figure 8 for target trials and supplementary Figure S6 for distractor trials. Furthermore, we conducted a comparison of correlations between distinct groups, healthy individuals and pwMS(BZDn), in different conditions (0-back, 1-back, 2-back). Our analysis revealed a significant dissimilarity in the correlations between these two study cohorts with a stronger positive correlation in HC subjects (p-value(0-back) = 0.0009, p-value(1-back) = 0.0015, p-value(2-back) = 0.0005). We did not observe an association between the 1/f slope modulation and the other cognitive scores: SDMT, VGLT and COWAT.

**Figure 8.**
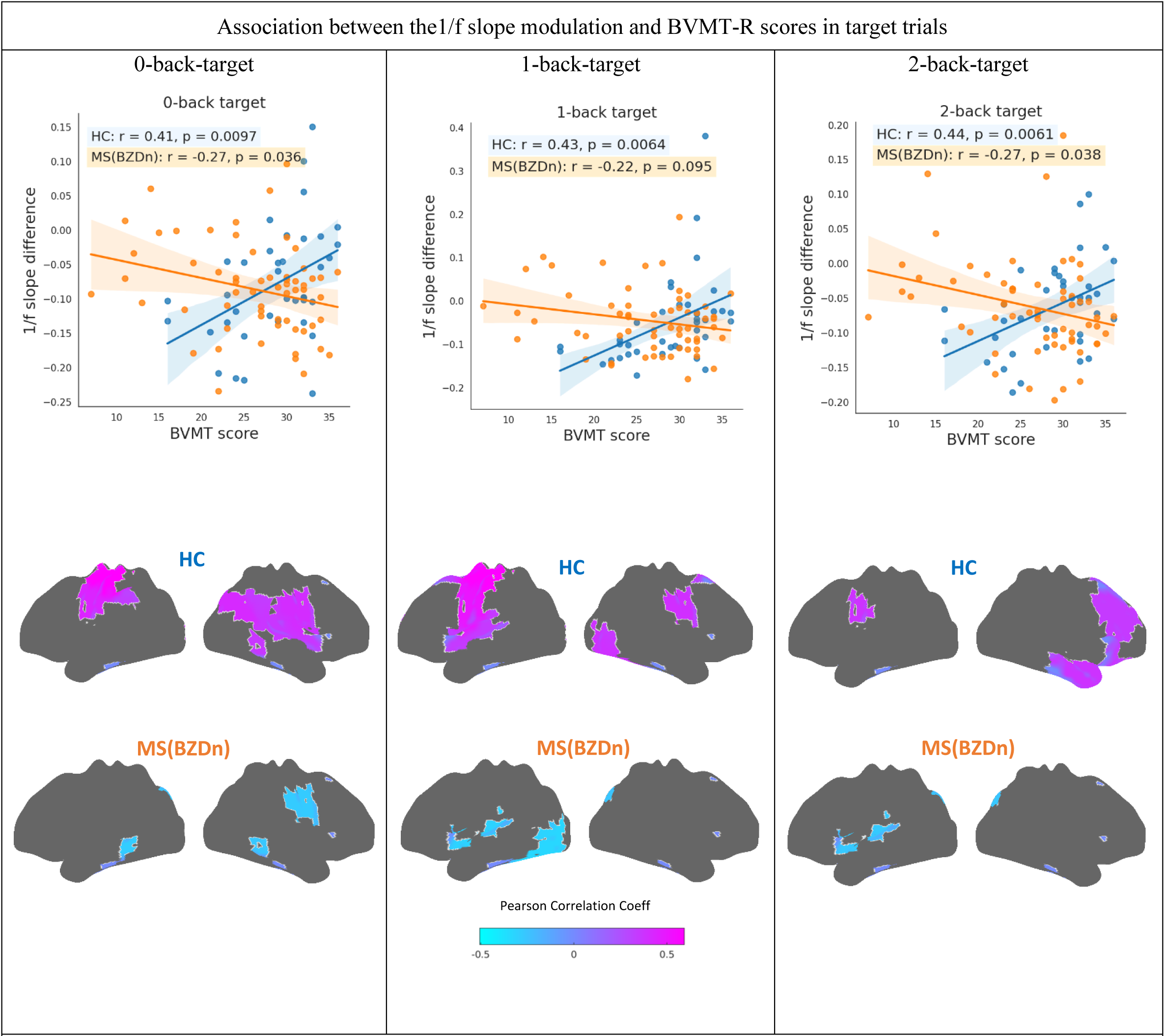
Correlation analysis between the 1/f slope modulation and BVMT-R scores both for averaged over the whole brain and in parcel level for the HC and pwMS(BZDn) groups in three conditions (0-back, 1-back and 2-back) in target trials. Only the parcels with significant correlations are shown.

## 4. Discussion

In the present study, we investigated stimulus-related modulation in the 1/f spectral slope of the brain during a visual-verbal working memory task. Our findings shed light on the interplay among neural activity, cognitive functions, and task performance.

### Steeper 1/f slope post stimulus

One of the primary observations of our study was the significant increase in the 1/f spectral slope following the onset of target stimuli. This suggests a profound impact of visual-verbal WM tasks with three cognitive load levels on the brain’s spectral dynamics. Similarly, Gyurkovics et al (Gyurkovics et al., 2022) found a steeper 1/f slope after auditory stimuli. This could suggest either a disruption of ongoing excitatory activity or an increase in inhibition level proportional to processing demands. It is important to note that the increase in 1/f slope is strongest in the sensorimotor cortex following target stimuli. To rule out the potential influence of button presses on 1/f slope changes in target trials, we also examined the 1/f slope change following distractor stimulus onset. Similarly, the 1/f slope became steeper after the distractor stimulus. This alteration in slope potentially indicates a reduction in the E/I ratio following the stimulus. Yet, the same effect arises across different brain regions following distractor trials. The 1/f changes can thus not solely be attributed to the requirement to push a button.

### Steeper post-distractor stimulus 1/f slope in HCs vs pwMS

Our study also revealed a difference in 1/f slope between HCs and pwMS, specifically after distractor stimulus presentation. The flatter 1/f slope after distractor trials in pwMS suggests a less pronounced inhibition of irrelevant information compared to HCs, highlighting a potentially reduced cognitive control in MS. This finding aligns with a previous study (Huiskamp et al., 2022) who used a large post-mortem MS dataset and a computational model to demonstrate that predominant inhibitory synaptic loss leads to network disinhibition and cognitive impairment in MS. Remarkably, the significantly flatter 1/f slope in the left inferior dorsal prefrontal cortex of pwMS emerged as a shared region in both 1-back and 2-back conditions. This particular brain parcel is known for its key role in motor planning and the maintenance of sustained attention and working memory, and executive functions.

### Correlation between 1/f slope and reaction time

A compelling finding from our study is the significant positive correlation between 1/f slope and reaction time both in the pre- and post-stimulus time window, particularly pronounced in the 2-back condition. Our results corroborate the findings of a recent study (Kałamała et al., 2023) which demonstrated that the 1/f slope is positively associated with adjusted reaction times as a behavioral performance. Moreover, the stronger correlation in the 1-back and 2-back conditions supports the involvement of more demanding WM processes, such as information updating and manipulation, which may contribute to the observed stronger relationship compared to the 0-back condition.

### The 1/f slope modulation and cognitive scores

Our study highlighted the clinical relevance of task-induced 1/f slope modulation by demonstrating significant positive and negative correlations with offline visuospatial memory as measured by BVMT-R for HCs and pwMS(BZDn). Our data show that healthy subjects with higher visuospatial performance tend to exhibit smaller modulation in the 1/f slope after stimulus onset, while in pwMS(BZDn), better performance on visuospatial tasks is associated with larger modulation in the 1/f slope. These significant correlations were observed in brain parcels within the parietal, temporal, and frontal lobes in HCs and parietal, temporal, and occipital lobes in pwMS(BZDn). These findings indicate the potential utility of task-induced modulation in the 1/f slope as a neurophysiological marker for assessing cognitive function in clinical contexts.

### The association between 1/f slope and alpha oscillations

Finally, we observed a significant positive correlation between alpha oscillatory power and the 1/f slope before and after stimulus presentation, indicating that higher alpha power is associated with a steeper 1/f slope. Interestingly, the changes in alpha power and 1/f slope post-stimulus onset vs. pre-stimulus occurred in opposite directions. While we found a steeper 1/f slope after stimulus onset, the expected post-stimulus decreases in alpha oscillatory activity (commonly referred to as alpha suppression (Feng et al., 2017; Foxe and Snyder, 2011)) were also observed; see Supplementary materials. This observation implies that oscillatory and non-oscillatory spectral components respond to stimuli independently.

## 5. Study limitations

Our study has a possible limitation related to the constrained time frame before and after the stimulus onset. Due to the nature of our task paradigm, we had to restrict the analysis to a one-second time window which lowered the frequency resolution to 1 Hz. Nonetheless, our dataset exhibited a high degree of accuracy in fitting the 1/f aperiodic component; see Supplementary Figure S7. It may prove beneficial to consider a broader time window before and after the stimulus onset in future research. Further, we arbitrarily determined the window size; other analyses enable the analysis of neurophysiological data in an intrinsically dynamic way (see e.g., (Rossi et al., 2023)). Finally, it is important to note that the 1/f slope has been suggested as a proxy for the E/I ratio, but alternative explanations have been proposed, e.g., the 1/f scaling is hypothesized as the result of the rate fluctuations in the cortical up and down states transition in the brain (Baranauskas et al., 2012) and also as a consequence arising from the damping of harmonic oscillators (Muthukumaraswamy and Liley, 2018).

## 6. Conclusion

In conclusion, our study revealed an increase in the steepness of the 1/f slope of the neuronal power spectra following a visual-verbal stimulation, and this observed broadband 1/f slope change is independent of the known alpha suppression. Furthermore, our results, indicating a more pronounced increase in 1/f slope in healthy subjects following the distractor stimuli compared to pwMS and also the association of 1/f slope modulation with behavioral and visuospatial working memory performance, underscore the potential significance of the 1/f spectral slope as a valuable proxy of cognitive functioning in MS.

## Acknowledgment

We would like to thank all participants for their time, enthusiasm, and commitment to participate.

## Data availability

Data is available upon reasonable request from the corresponding author.

## Supplementary materials

Stimulus-related modulation in the 1/f spectral slope suggests an impaired inhibition of irrelevant information in people with multiple sclerosis. Fahime Akbarian et al.

**Figure S1.**
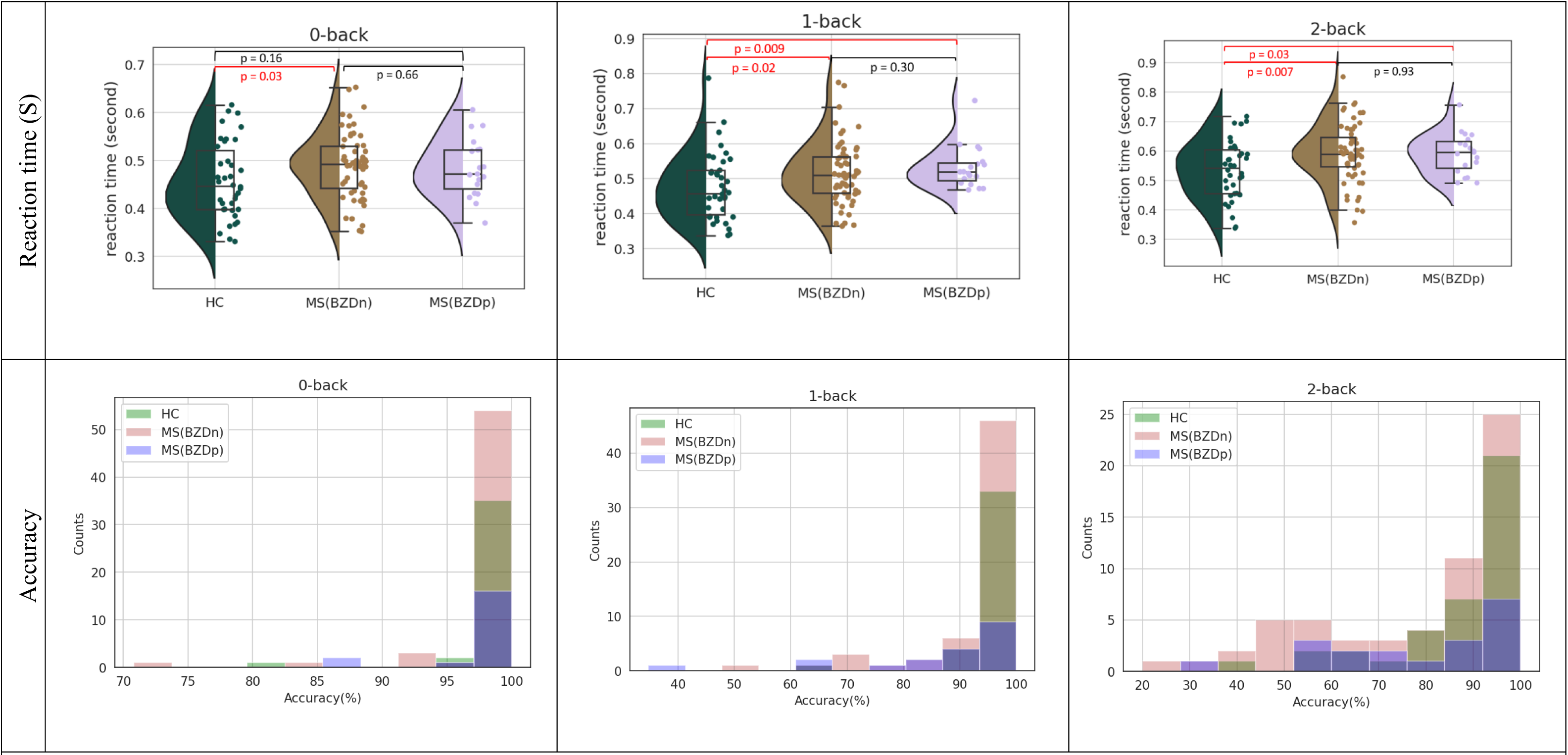
Reaction times and accuracy. The response accuracy is defined as the number of correct answers over the total target trials for the specific task condition. Wilcoxon rank-sum test and Mann-Whitney U test were used to compare the reaction times and accuracy respectively.

**Figure S2.**
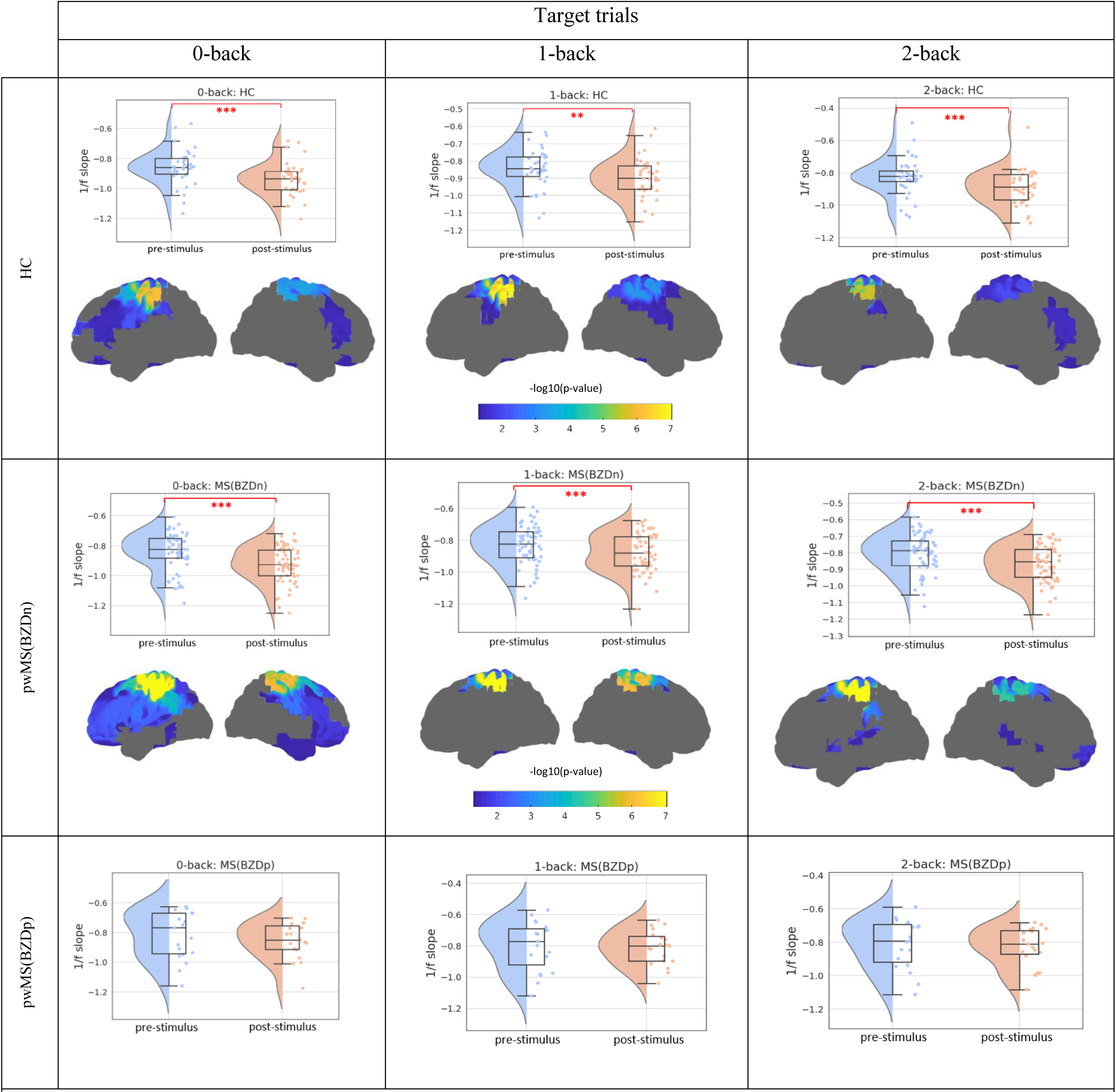
The pre- vs post-stimuli 1/f slope in target trials and three conditions (0-back, 1-back and 2-back), for both the 1/f slope averaged over the whole brain and at parcel level. Note that the -log10(p-value)>1.3 indicated the significant values and that p-values have been corrected for multiple comparisons across the 42 parcels. The Wilcoxon signed-rank test was used to compare the pre- and post-stimuli 1/f slope. First row: HC subjects, second row: pwMS(BZDn), third row: pwMS(BZDp). *:0.001<p<0.05, **: 0.0001<p<0.001, ***: 0.00001<p<0.0001.

**Figure S3.**
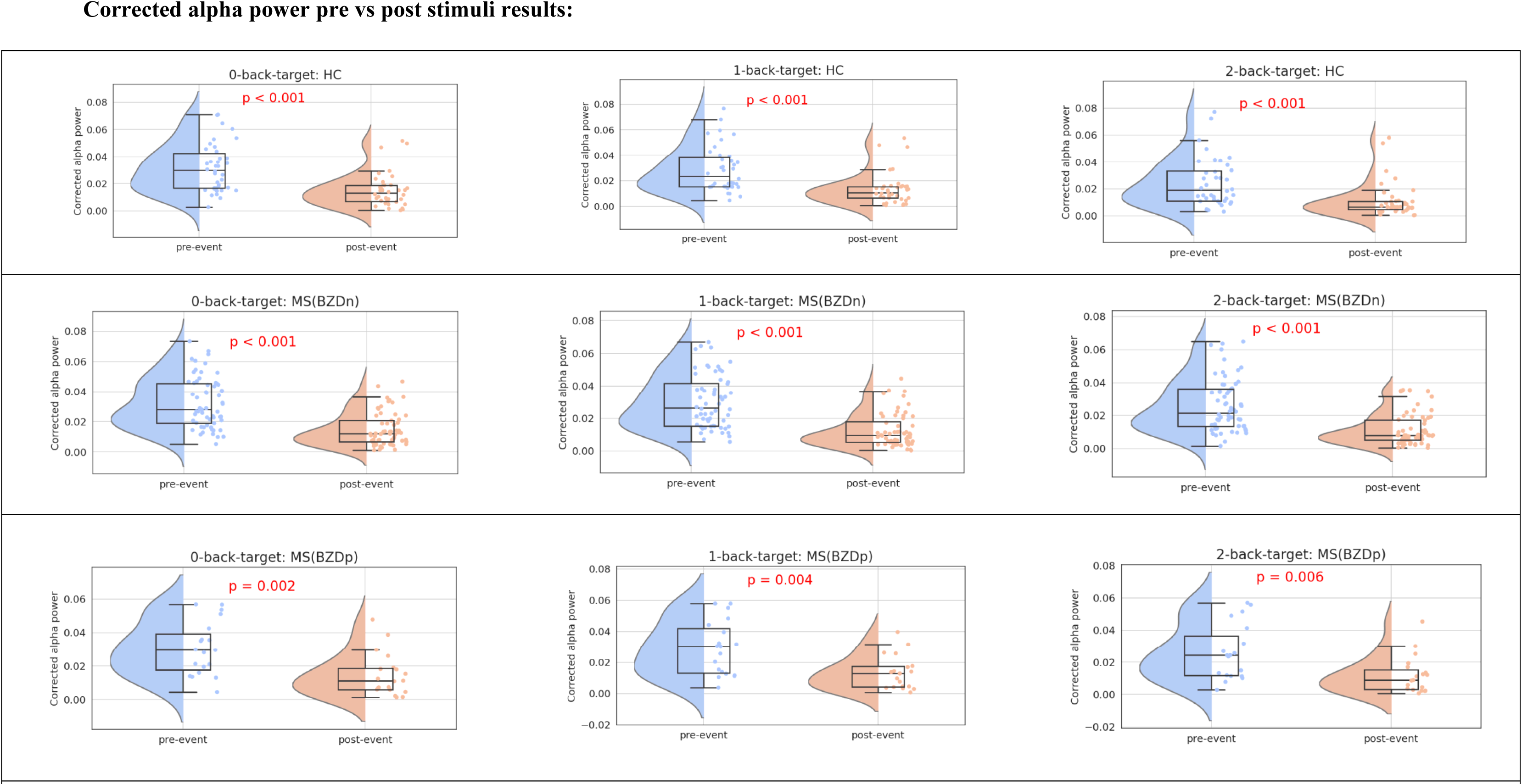
The pre- vs post-stimuli **corrected alpha power** in **target trials** and three conditions (0-back, 1-back and 2-back) for corrected alpha power averaged over the whole brain. The Wilcoxon signed-rank test was used to compare the pre- and post-stimuli corrected alpha power. First row: HC subjects, second row: pwMS(BZDn), third row: pwMS(BZDp).

**Figure S4.**
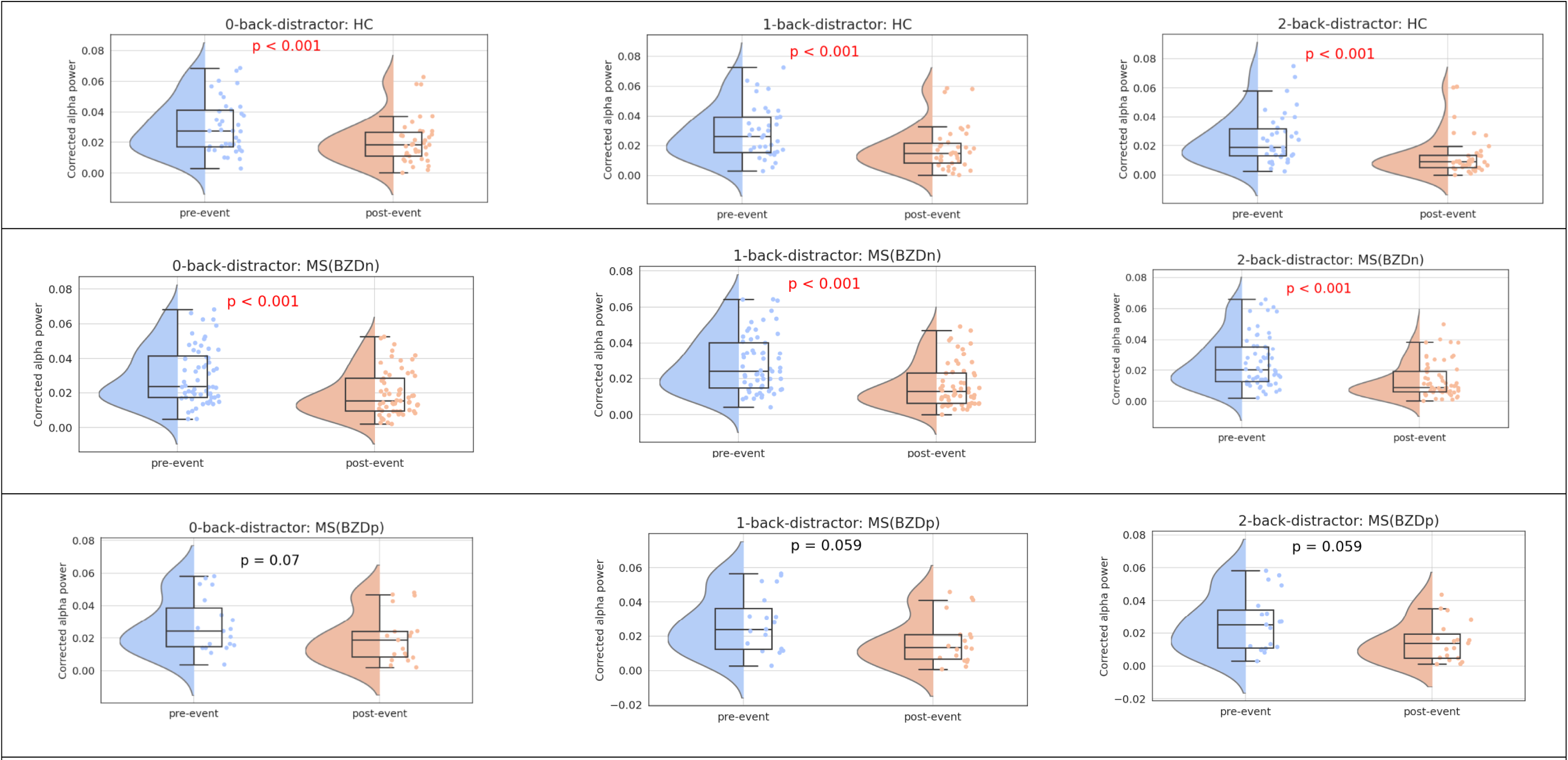
The pre- vs post-stimuli **corrected alpha power** in **distracor trials** and three conditions (0-back, 1-back and 2-back) for corrected alpha power averaged over the whole brain. The Wilcoxon signed-rank test was used to compare the pre- and post-stimuli corrected alpha power. First row: HC subjects, second row: pwMS(BZDn), third row: pwMS(BZDp).

**Figure S5.**
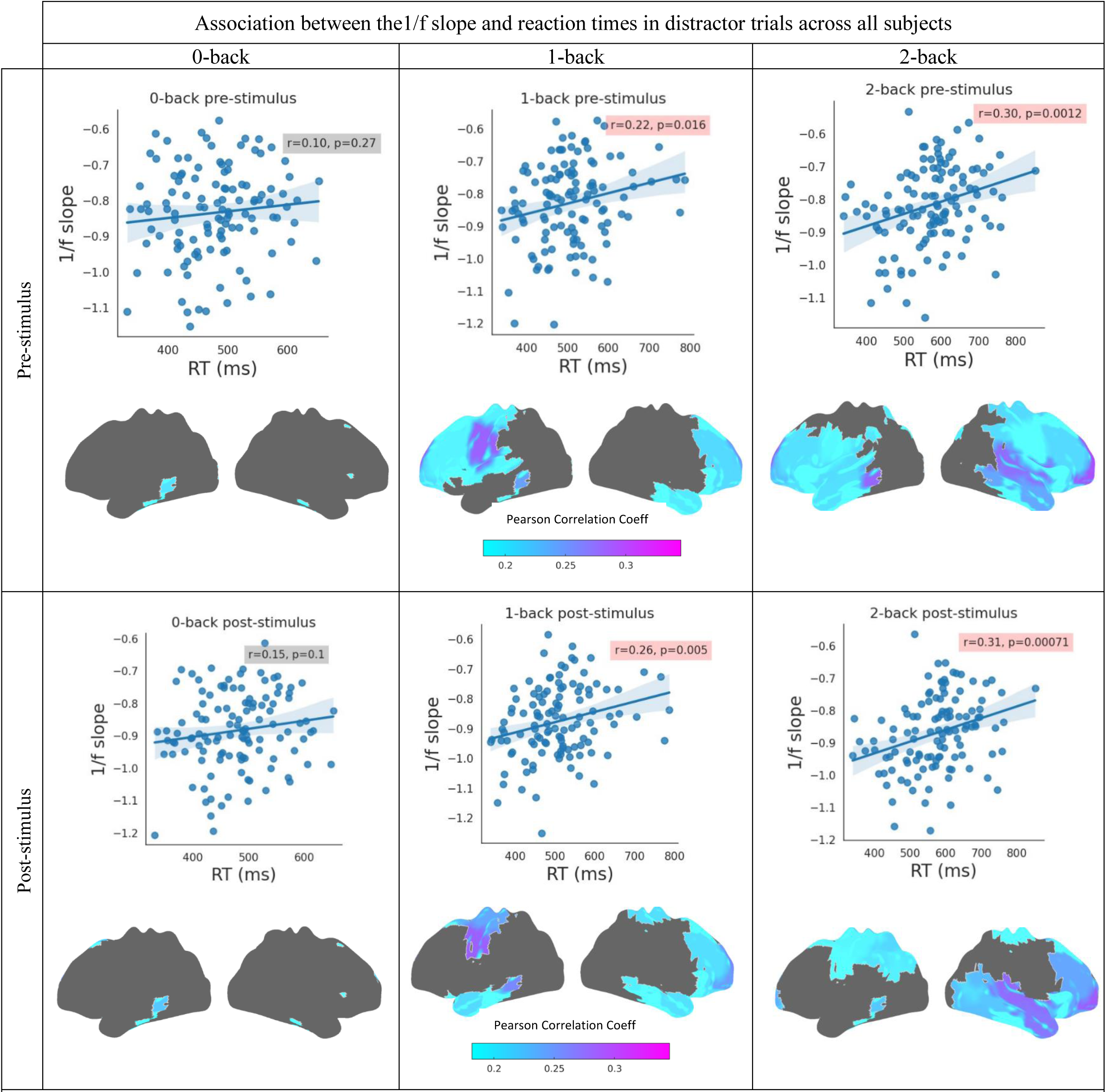
Correlation analysis between reaction time and 1/f slope of the time window one second pre- and one second post-distractor stimulus onset both for averaged over whole brain and in parcel level over all subjects. In each grid, the scatter plot shows the correlation for the averaged 1/f slope over whole brain and reaction times, and the cortical map shows the parcels with significant correlation for the parcel level analysis. We included all subjects to increase the statistical power.

**Figure S6.**
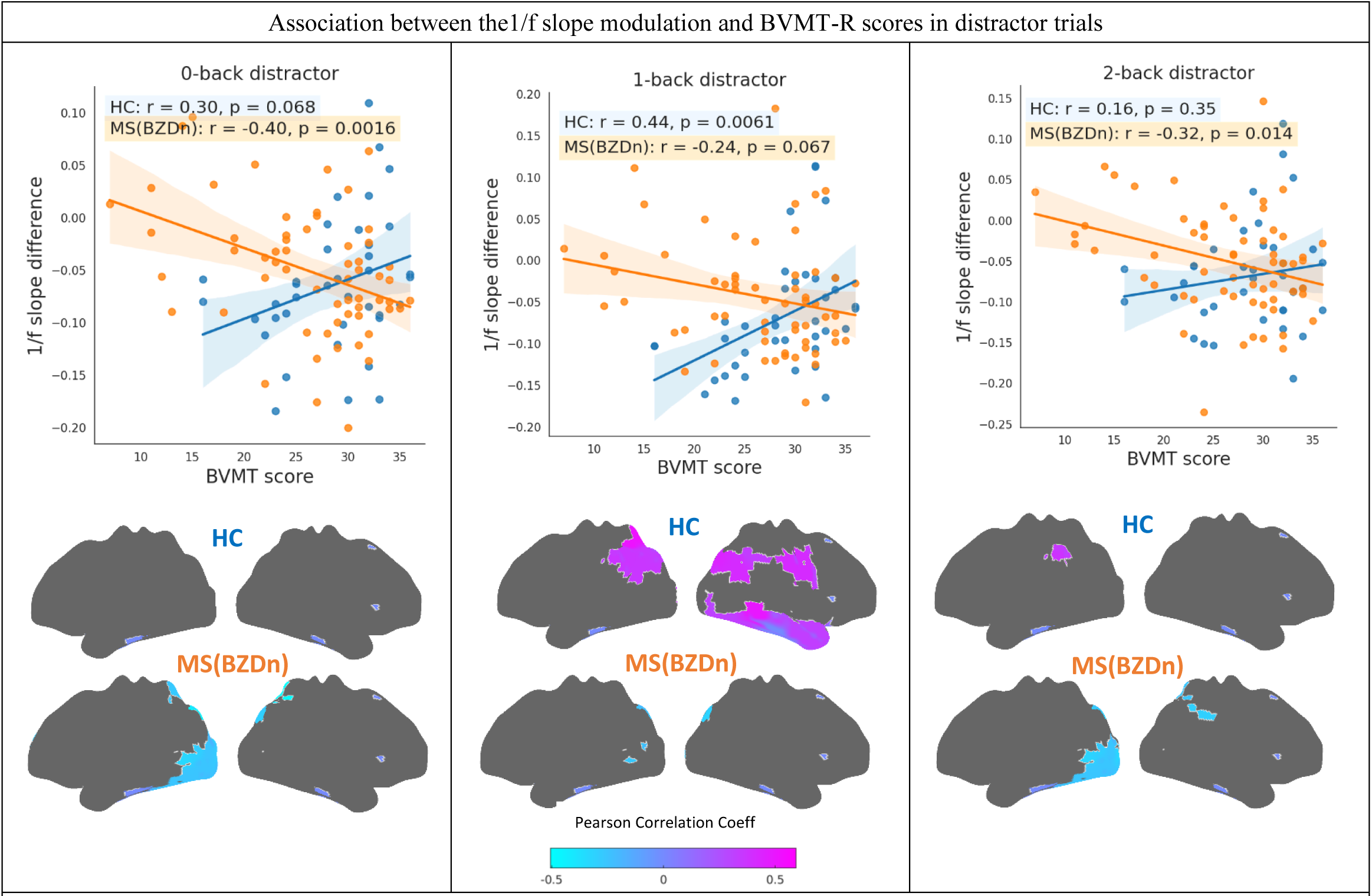
Correlation analysis between the 1/f slope modulation and BVMT-R scores both for averaged over the whole brain and at parcel level for the HC and pwMS(BZDn) groups in three conditions (0-back, 1-back and 2-back) in distractor trials. Only the parcels with significant correlations are shown.

**Figure S7.**
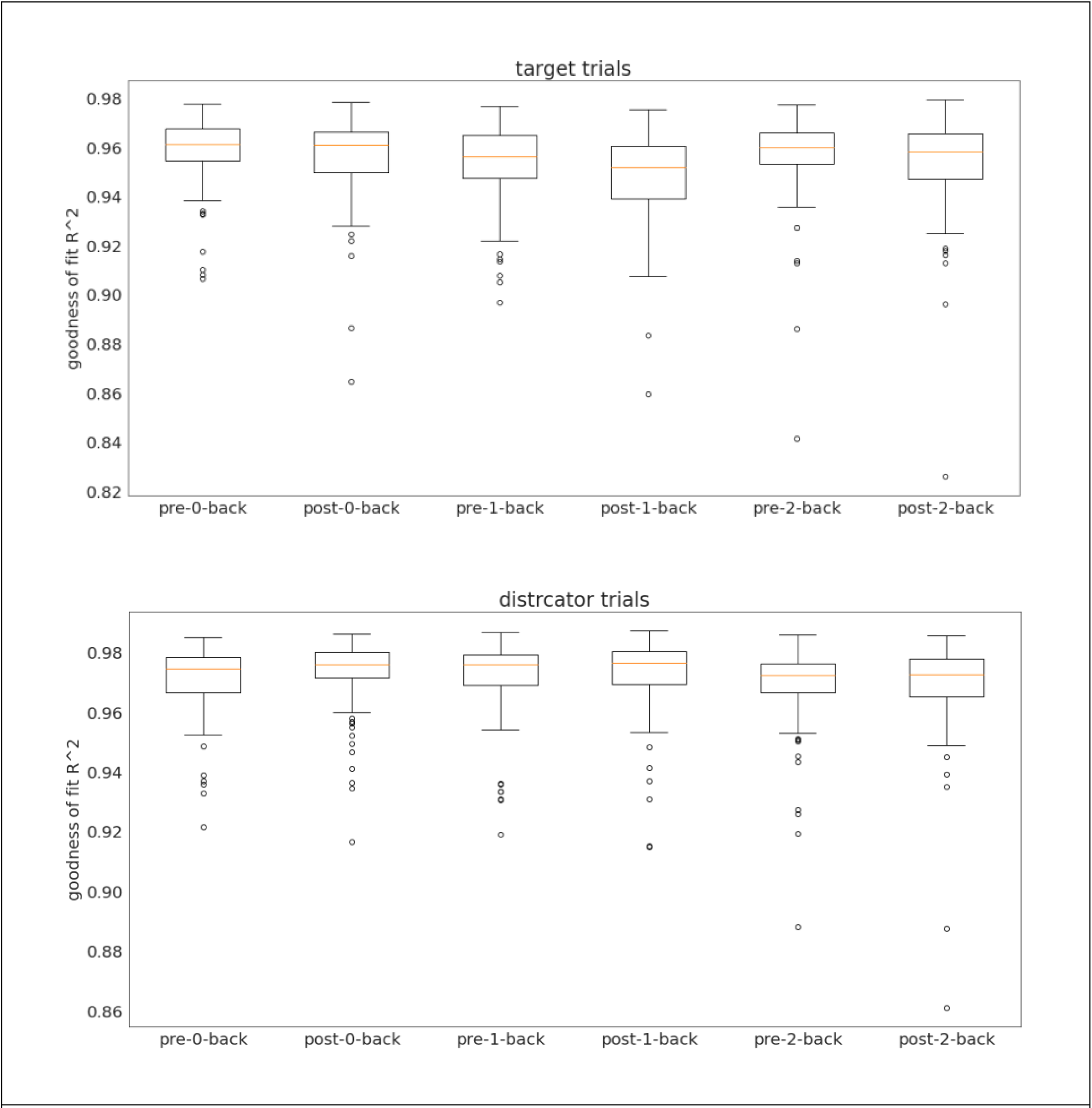
Goodness of Fit. Box plots of the R^2^ parameter of FOOOF fitting of power spectrum densities within pre- and post-stimulus time windows for each WM load condition for all subjects in target (top) and distractor trials (bottom).

**Table S1.** List of regions used in the parcel level analyses.

|  | ROI |
| --- | --- |
| 'ROI 1' | 'L Cuneus' |
| 'ROI 2' | 'R Cuneus' |
| 'ROI 3' | 'L Inf Occ' |
| 'ROI 4' | 'R Inf Occ' |
| 'ROI 5' | 'L Supramarginal' |
| 'ROI 6' | 'R Supramarginal' |
| 'ROI 7' | 'L Sup Temp' |
| 'ROI 8' | 'R Sup Temp' |
| 'ROI 9' | 'L Lat SMC' |
| 'ROI 10' | 'R Lat SMC' |
| 'ROI 11' | 'L Sup Parietal' |
| 'ROI 12' | 'R Sup Parietal' |
| 'ROI 13' | 'L Middle Occ' |
| 'ROI 14' | 'R Middle Occ' |
| 'ROI 15' | 'L Sup Occ' |
| 'ROI 16' | 'R Sup Occ' |
| 'ROI 17' | 'L Ant Temp' |
| 'ROI 18' | 'R Ant Temp' |
| 'ROI 19' | 'L Medial SMC' |
| 'ROI 20' | 'R Medial SMC' |
| 'ROI 21' | 'L Angular' |
| 'ROI 22' | 'R Angular' |
| 'ROI 23' | 'L VL PFC' |
| 'ROI 24' | 'R VL PFC' |
| 'ROI 25' | 'L Occ pole' |
| 'ROI 26' | 'R Occ pole' |
| 'ROI 27' | 'L Sup PFC' |
| 'ROI 28' | 'R Sup PFC' |
| 'ROI 29' | 'L Sup Dorsal PFC' |
| 'ROI 30' | 'R Sup Dorsal PFC' |
| 'ROI 31' | 'L Orbitofrontal' |
| 'ROI 32' | 'R Orbitofrontal' |
| 'ROI 33' | 'L Post Temp' |
| 'ROI 34' | 'R Post Temp' |
| 'ROI 35' | 'L Inf Dorsal PFC' |
| 'ROI 36' | 'R Inf Dorsal PFC' |
| 'ROI 37' | 'Medial PFC' |
| 'ROI 38' | 'Post Precuneus' |
| 'ROI 39' | 'Posterior Cingulate Cortex' |
| 'ROI 40' | 'Ant Precuneus' |
| 'ROI 41' | 'L Inf Parietal' |
| 'ROI42' | 'R Inf Parietal' |

